# Supervised Learning Model Predicts Protein Adsorption to Carbon Nanotubes

**DOI:** 10.1101/2021.06.19.449132

**Authors:** Nicholas Ouassil, Rebecca L. Pinals, Jackson Travis Del Bonis-O’Donnell, Jeffrey Wang, Markita P. Landry

## Abstract

Engineered nanoparticles are advantageous for numerous biotechnology applications, including biomolecular sensing and delivery. However, testing the compatibility and function of nanotechnologies in biological systems requires a heuristic approach, where unpredictable biofouling via protein corona formation often prevents effective implementation. Moreover, rational design of biomolecule-nanoparticle conjugates requires prior knowledge of such interactions or extensive experimental testing. Toward better applying engineered nanoparticles in biological systems, herein, we develop a random forest classifier (RFC) trained with proteomic mass spectrometry data that identifies proteins that adsorb to nanoparticles, based solely on the protein’s amino acid sequence. We model proteins that populate the corona of a single-walled carbon nanotube (SWCNT)-based optical nanosensor and study whether there is a relationship between the protein’s amino acid-based properties and the protein’s adsorption to SWCNTs. We optimize the classifier and characterize the classifier performance against other models. To evaluate the predictive power of our model, we apply the classifier to rapidly identify proteins with high binding affinity to SWCNTs, followed by experimental validation. We further determine protein features associated with increased likelihood of SWCNT binding: high content of solvent-exposed glycine residues and non-secondary structure-associated amino acids. Conversely, proteins with high content of leucine residues and beta-sheet-associated amino acids are less likely to form the SWCNT protein corona. The classifier presented herein provides a step toward undertaking the otherwise intractable problem of predicting protein-nanoparticle interactions, which is needed for more rapid and effective translation of nanobiotechnologies from *in vitro* synthesis to *in vivo* use.

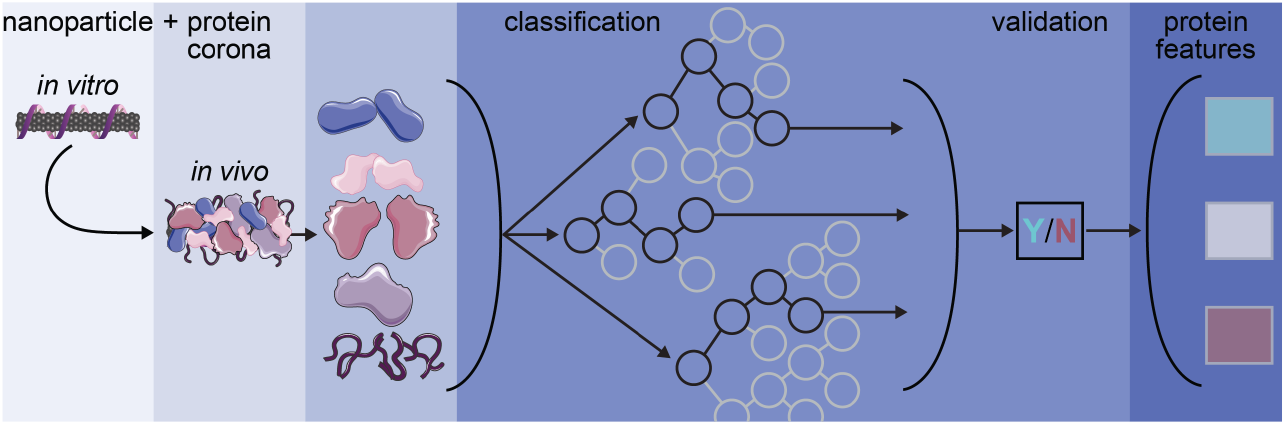

## Introduction

Engineered nanoparticles are poised to transform how we undertake biological sensing,^1–3^ imaging,^4–6^ and delivery:^7–9^ nanoscale materials enable localization within otherwise inaccessible biological environments and exhibit highly tunable physicochemical properties to tailor function. Different nanoparticle platforms offer application-dependent advantages, such as near-infrared fluorescent nanoparticles for through-tissue imaging^10,11^ or biodegradable nanoparticles for *in vivo* delivery.^12–14^ In particular, single-walled carbon nanotubes (SWCNTs) are well-suited for biological sensing and imaging due to their tissue-transparent and photostable near-infrared fluorescence, in addition to their readily modifiable surface.^15–17^ As such, SWCNTs have been functionalized with biomolecules including single-stranded DNA to create neurotransmitter nanosensors,^18–20^ with peptide mimetics to form protein nanosensors,^21^ and with proteins to construct viral nanosensors.^22^ Similarly, the large SWCNT surface area enables cargo attachment such that SWCNTs can be loaded with DNA plasmids or small interfering RNAs, translocating these functional biomolecules into cells for gene expression and silencing applications.^23–26^ Optimizing these biomolecule-nanoparticle interactions is key in enhancing nanotechnology function, and a deeper understanding of these interfacial interactions would enable more rational conjugate designs. As such, the capability to predict nano-bio interactions would aid the design phase of nanobiotechnologies by lessening the need to experimentally test innate interactions of each biomolecule with each nanoparticle of interest.

Although such aforementioned nano-bio interactions are required for function, conversely, biofouling of nanobiotechnologies resulting from undesired nano-bio interactions often inhibits intended nanoparticle function. SWCNTs and other nanotechnologies more broadly suffer from as-of-yet unpredictable interactions with the biological environments in which they are applied. When engineered nanoparticles are introduced into biological systems, endogenous proteins rapidly bind to the nanoparticle surface.^27–29^ This phenomenon is known as protein corona formation. Protein adsorption often decreases the ability of the nanoparticle to interact with its surrounding environment, such as sensing nearby analytes^30–32^ or navigating biological barriers.^33–35^ Due to its inherent complexity, the protein corona remains a poorly understood phenomenon limiting the efficiency with which nanoparticle-based technologies are applied in biological systems.^33,36,37^ Limitations in our understanding of corona formation arise from a convolution of diverse nanoparticle properties (dominated by surface characteristics) and the complexity of biological environments.^28,34,38,39^ Yet, knowledge of the proteins adsorbed in this corona phase would enable better prediction of the biological identity, and thus fate, of the applied nanotechnologies.^40,41^ Experimental testing to fully characterize the protein corona on all synthesized nanoparticle constructs within all intended biological environments is laborious and costly: while recent work has made headway toward high-throughput experimental methods,^42,43^ the most common strategies rely on labor-intensive mass spectrometry-based proteomics.^38,44,45^ The ability to predict the protein corona that will form on nanoparticles *in vivo* remains a challenge that, if overcome, would improve applied nanotechnology performance.

Pattern recognition techniques, including those of machine learning, offer a route to characterize protein-nanoparticle interactions in a high-throughput manner across this extensive design space of nanoparticles applied in different biological systems. Previous work pioneering this idea applied random forest classification to predict what proteins adsorb to silver nanoparticles in biologically relevant environments,^44^ and has been expanded to larger nanoparticle libraries.^46^ However, certain aspects stand to be refined, such as setting the threshold of whether a protein is classified as in or out of the corona. Other work has examined protein-nanoparticle complexes using a fluorometric assay to guide prediction of corona formation, though issues arise in characterizing fluorophore interactions on graphene-based substrates.^47^ More broadly, most predictive modeling efforts involving nanoparticles applied in biology consider cellular- or organism-level responses, such as cellular association,^45,48^ toxicity,^49^ *in vivo* fate,^41^ and therapeutic efficacy.^48,50^ Toward protein-SWCNT conjugate design, some predictive modeling has informed protein candidates that exhibit natural affinity for the graphitic SWCNT surface.^23^ For example, Di Giosia *et al*. implemented a molecular docking model to determine a panel of proteins that interact with the carbon nanotube surface.^51^ Yet, this strategy of predicting protein corona identity requires protein structural information and is low throughput, both computationally and in experimental validation. Our workflow expands on this body work by classifying protein attachment to SWCNTs based only on protein sequence, as well as redefining metrics for determining placement in-corona.

Herein, we develop a classifier to investigate the relationship between a protein’s amino acid sequence and a protein’s binding propensity to carbon nanotubes. Our purpose is two-fold: as one objective, we aim to predict which protein-SWCNT interactions to expect in biological environments. This knowledge will inform implementation of anti-biofouling strategies toward effective biological application of nanoparticles. Our second objective is to predict high-affinity protein binders to SWCNTs and protein features associated with such binding affinity, to improve the process of protein-nanoparticle construct design.^23^ Toward these ends, we build and optimize a random forest classifier applied to protein adsorption on SWCNTs. We relate protein properties (derived from protein sequence data) to whether proteins are in or out of the corona phase on SWCNTs (experimentally determined by quantitative mass spectrometry-based proteomics). Specifically, we focus on protein corona formation on (GT)_15_-SWCNTs due to their demonstrated applicability for dopamine sensing,^18,19^ however, the workflow is generalizable to other nanoparticles. We train our classifier using mass spectrometry-based proteomic data characterizing the corona formed on (GT)_15_-SWCNTs in two relevant bioenvironments: the intravenous environment (blood plasma) and the brain environment (cerebrospinal fluid).^52^ We find that our classifier can precisely target the small number of proteins that adsorb to our nanoparticle. Furthermore, we identify population distribution changes among the most important protein properties to gain insight on how our classifier identifies positive targets. Namely, high content of glycine residues (particularly solvent-exposed residues) and amino acids not associated with secondary structure domains (alpha helix, beta sheet, and turns) lead to favorable SWCNT binding, whereas high content of leucine residues and amino acids associated with planar beta-sheet domains lead to unfavorable SWCNT binding. Finally, we test our model with an entirely new set of proteins and perform quantitative protein adsorption experiments to validate the model’s in vs. out of corona predictions.^32^ Our results suggest that this classifier can serve as a tool to understand how protein sequence influences nanotube binding.

## Results and Discussion

### Experimentally determined protein corona composition on (GT)_15_-SWCNTs

The protein corona dataset was experimentally generated from a selective adsorption assay that quantified protein amounts present on nanoparticles via liquid chromatography tandem mass spectrometry (LC-MS/MS) characterization.^52^ With this assay, corona proteins were determined for (GT)_15_-SWCNTs incubated in either human blood plasma or cerebrospinal fluid (CSF) of the brain (attached datasheet, reproduced from ^52^). The absolute protein abundance and relative enrichment or depletion (compared to the control sample of the biofluid alone) were used to indicate whether a particular protein was considered to be in the corona, as will be described in a later section. We included four protein corona datasets: (GT)_15_-SWCNTs in blood plasma, (GT)_15_-SWCNTs in cerebrospinal fluid, the total set with biofluid labels, and the total set naïve of biofluid labels. The biofluid label refers to the knowledge of where the protein originated (blood plasma or CSF).

### Protein property database development from protein sequence

We next curated a protein property database to use with our classifier. We used the amino acid sequence of each protein from the annotated protein database, UniProt,^53^ to construct an array of predicted physicochemical protein properties with the BioPython package (**Table S1**; see Database Development section in **Methods**).^54^ UniProt also provides biological protein properties such as gene ontology, sequence annotations, and specific functional regions, therefore we compared how the inclusion of these other properties influenced classifier performance (**Figure S1**). However, our final classifier was based solely on amino acid sequence data to avoid potential issues of less well-studied proteins that have no empirically derived properties and/or no annotated features. Developing a database in this manner expands the number of possible proteins that can be tested because the classifier does not require prior information on the annotated protein sequence, nor on interactions between the protein and nanoparticle. Thus, requiring access to only the protein’s amino acid sequence enables facile expansion to model new experimental datasets or to select new nanoparticle-binding proteins of interest.

The amino acid sequence of a protein provides valuable information including the percentage of a specific amino acid within the full protein; however, spatial organization is disregarded. To complement the sequence-derived dataset, we added the parameter of solvent accessibility that estimates the exposed protein surface area. We implemented NetSurfP 2.0^55^ to predict the number of exposed residues of a particular protein using the amino acid sequence, normalized by either the total number of amino acids or the total number of exposed amino acids. These two choices of normalization provide information on amino acid content on the surface relative to the full protein or relative to only other surface-exposed residues, respectively.

### Thresholding to determine protein placement: in or out of the corona

The decision of whether a protein was categorized as in or out of the corona was made using the protein abundance data from LC-MS/MS experiments. Proteins were placed into the corona based on two criteria: (i) relative change and (ii) an abundance threshold. First, if a protein was more abundant in the nanoparticle-bound case than it was in the control solution of the native biofluid without any nanoparticles present (i.e., enrichment on the nanoparticle), then it was classified as in the corona. Second, the remaining proteins were ordered by abundance and fit to an exponential distribution. Increasing the power of the exponential leads to a higher in-corona threshold, placing fewer proteins in the corona. Importantly, this thresholding approach reflects that lower abundance of a protein in the corona relative to its abundance in the biofluid (i.e., depletion on the nanoparticle) does not necessarily mean that protein is out of the corona; a protein that is significantly depleted can still be present in the corona with a high absolute quantity. The thresholding method that we have developed is discussed further in the Methods section.

### Random forest classifier development and verification using established protein property database

We implemented a random forest classifier (RFC) to classify corona proteins on (GT)_15_-SWCNT nanoparticles. Although we focus on protein corona characterization with one nanoparticle type, SWCNTs, it is worthwhile to note that these classifiers do not require any information regarding the nanoparticle itself. We chose to pursue an RFC because this is an ensemble method with a well-known ability to be resistant to overfitting by employing several weak learners that fit to different parameters,^56^ Moreover, an RFC produces highly interpretable results. Implementing an RFC is in line with previously published work.^44,46^ An example tree to illustrate this process is provided as **Figure S2**. To confirm the choice of an RFC over other potential classifiers, we tested an assortment of classifier types (**Figure S3**). The highest performing classifiers were the RFC and a bagging classifier using decision trees, based on a sum of the metrics of accuracy, precision, and recall. We selected the RFC for this study because the precision (0.671) and accuracy (0.747) values were superior to that of the bagging tree, while retaining a similar area under the receiver operating curve (AUC; 0.700). AUC is a frequently used measure for understanding sensitivity and specificity of the classifier. Moreover, the RFC provided the highest precision (positive predictive value), which is favorable for the most straightforward application of classifier output for nanobiotechnology optimization. However, the bagging tree did perform better than the RFC in recall (bagging tree, 0.528; RFC 0.505).

Due to the imbalance in our LC-MS/MS experimental dataset (i.e., unequal number of proteins in either class), we up-sampled our minority class (in corona; ~30% in corona without up-sampling in the total dataset). This up-sampling ensures that each time the classifier was trained, we were able to recover an appropriate amount of the minority class. For this reason, the classifier was validated using a stratified shuffle split repeated 100 times. Moreover, we noticed that generalization of this classifier could be improved, especially when considering the recall was below 0.5. To address this issue, a synthetic minority over-sampling technique (SMOTE)^57^ was implemented to generate new “proteins” in the minority class (in corona). This analysis revealed that the most precise and accurate results for our classifier were obtained when the minority/majority ratio in SMOTE was 0.5/1.0 (**Figure S4**; precision 0.678 and accuracy 0.749). The recall of our classifier was improved marginally from 0.497 to 0.512. Introducing the described methods expanded the number of proteins that were placed in the corona. However, this SMOTE ratio offers a tunable handle: if an experimenter preferred higher recall values, a ratio of 1 provides a recall of 0.587, although reducing precision to 0.571.

Using an RFC, classification tests were run on the total naïve dataset of proteins marked as being in or out of the corona with the aforementioned thresholding method. The classifier performance was scored for a range of thresholding powers (**Figure 1a**). The classifier was then refreshed and the standard protocol for training the classifier was repeated to gather metrics related to classification: accuracy, area under the receiver operating curve (AUC), precision, and recall. The metrics were recorded until a thresholding power of 3.5, at which point higher powers considerably reduced the number of proteins counted in the plasma corona and many metrics drastically declined in their performance. We ultimately selected a power of 2.25 because this power provided the best compromise between accuracy (0.747), AUC (0.691), and precision (0.648), while only suffering slightly in recall (0.570). All reported results for the remainder of this work use a power of 2.25 for placing proteins in the nanoparticle corona.

**Figure 1.**
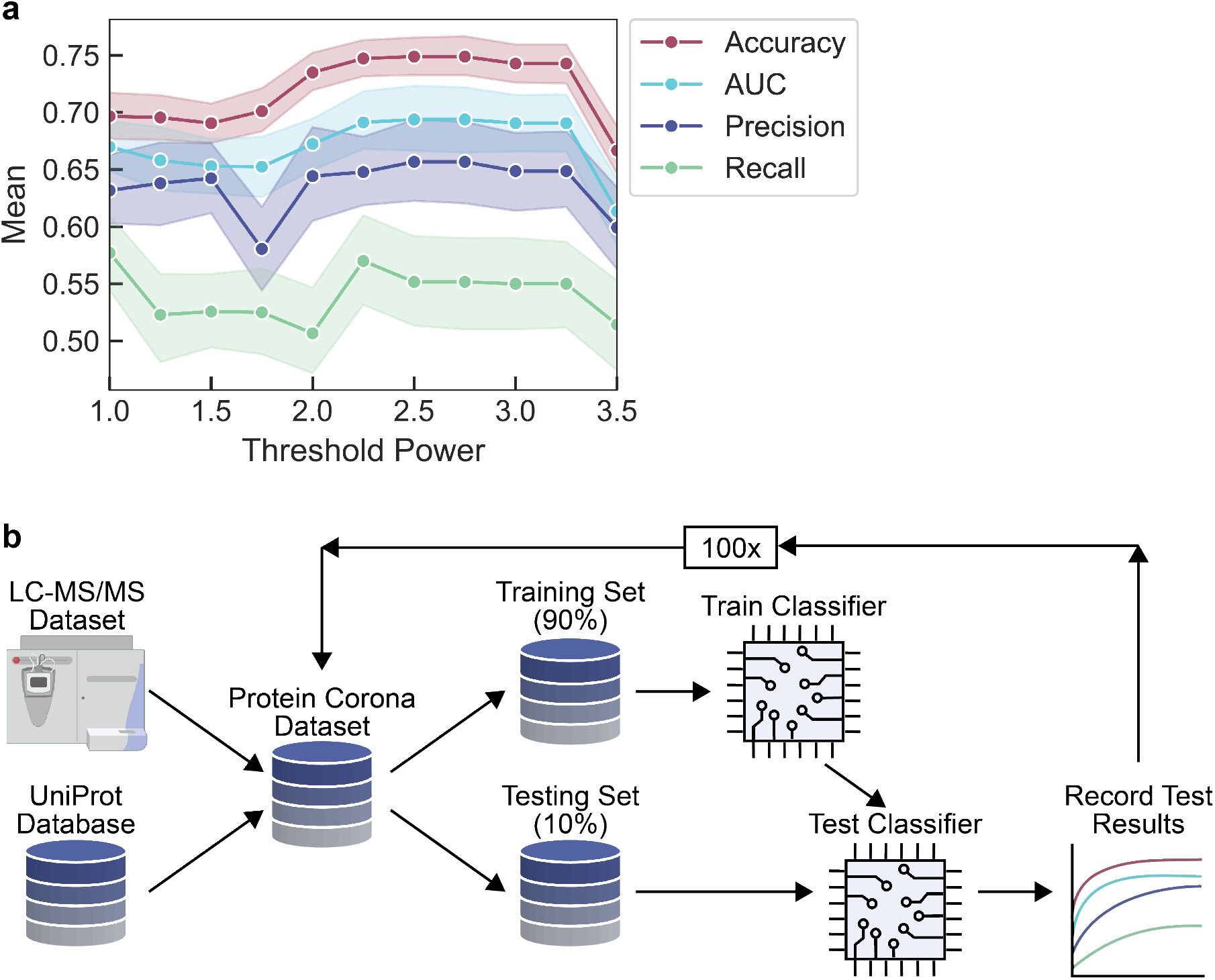
Random forest classifier (RFC) development and workflow for determining proteins in vs. out of the corona phase on (GT)_15_-SWCNTs. (a) Metrics of accuracy, area under the receiver operating curve (AUC), precision, and recall recorded as a function of threshold power for labeling proteins as in vs. out of the corona. A threshold power value of 2.25 is selected for subsequent analyses due to the optimal combination of the recorded metrics. Shaded error bars represent 95% confidence intervals. (b) RFC workflow used in splitting-based predictions. Liquid chromatography-tandem mass spectrometry (LC-MS/MS) experimentally provides protein corona composition. LC-MS/MS data is combined with protein properties derived from the protein sequence (UniProt database with BioPython package for analysis) to form a total dataset. The total dataset is split 90% into training data and 10% into test data. Training data is used to train a reset classifier then test data is used to score the trained classifier. Results are recorded and the process is repeated.

During development, stratified shuffle split validation was used to check the success of our classifier with respect to accuracy, AUC, recall, and precision. The dataset was divided into a training and test set at the beginning of each split, then the training set was fit to an untrained classifier. Next, predictions were made on the test set and compared with our true answers. The results from this classifier were saved and the process was repeated with the classifier naïve at the beginning of each iteration, as graphically depicted in **Figure 1b**. This method was used to ensure that the subset of proteins generated more accurate metrics for the classifier, considering each protein revolved into the test set during one of the folds. Statistics represented in this work were generated from the n trials used in this verification step.

The first trial was with two datasets, total set labeled vs. total set naïve (**Figure 2a**). The only difference between these two datasets was the inclusion of one Boolean column that dictates from which biofluid a protein originated. We observe that the inclusion of this “biofluid of origin” information does not improve the classification ability on our complete dataset. Thus, we deemed this column unnecessary to include for future runs. Moreover, keeping this column would have made our classifier less generic when selecting new proteins that may not be present in blood plasma or CSF.

**Figure 2.**
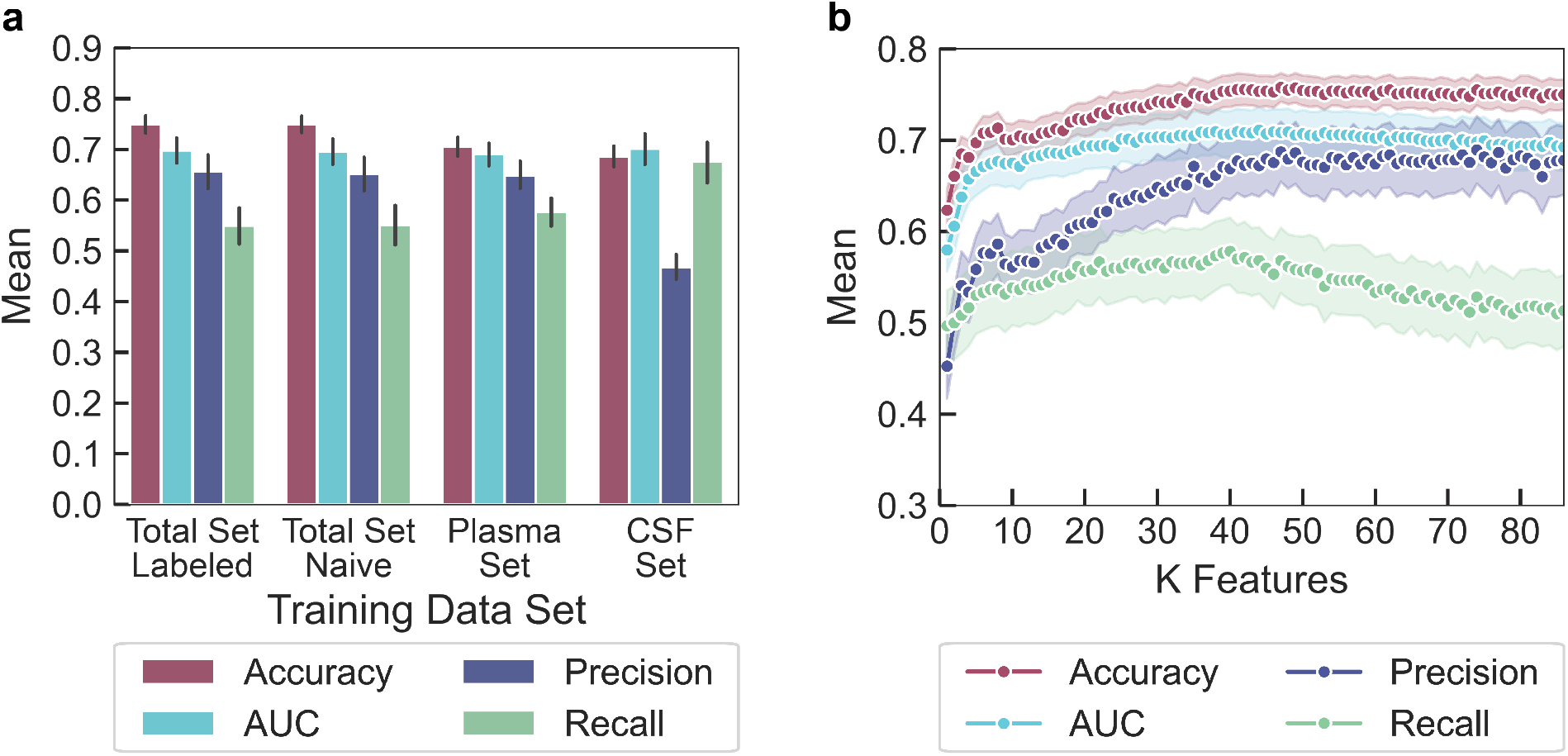
Classifier results on different training datasets and with varied feature inputs. (a) RFC trained on the full protein set (with or without the label of origin biofluid) or each individual biofluid (plasma or CSF). Negligible differences arise between the RFC’s ability to classify the total set with or without the biofluid label (total set labeled compared to total set naïve), denoting that the biofluid label feature does not resolve the cross-fluid classification problem. Training the RFC on one biofluid and testing against the second biofluid produces similar metrics except for precision, attributable to a few proteins labeled in the corona of one biofluid but not the other. Error bars represent 95% confidence intervals. (b) RFC trained on the total naïve protein corona set, with features sorted by ANOVA and added to the classifier from highest to lowest importance. At approximately 40 features, classification ability begins to plateau for all metrics except recall. By 89 features, there is a decline in recall but marginally enhanced precision. Shaded error bars represent 95% confidence intervals.

We next trained the classifier on corona proteins present in one biofluid and attempted to predict corona proteins present in another biofluid. For this case, instead of splitting the data 90% training/10% testing, the classifier was trained on one complete dataset, then a subset of the second dataset was used as the testing set. We repeated this approach 100 times to generate statistics for the classifier. We notice similar results in AUC (CSF: 0.702, plasma: 0.691) and accuracy (CSF: 0.687, plasma: 0.706) independent of which biofluid the classifier was trained on (**Figure 2a**). However, there is a difference in precision (CSF: 0.469, plasma: 0.649) and recall (CSF: 0.676, plasma: 0.577) for each of these classifiers, arising from the inclusion of a few of proteins that are present in the corona formed on (GT)_15_-SWCNTs from one biofluid and are not present in the corona formed on (GT)_15_-SWCNTs from the other biofluid (e.g., serotransferrin found in the CSF corona and haptoglobin found in the plasma corona). This discrepancy occurs because our classifier has no context of which proteins are in the corona formed from which biofluid, and thus there is no method of adjusting for proteins portraying contradicting adsorptive behavior across biofluids. However, this classification discrepancy only occurs for a few proteins (13 proteins out of 38 duplicate proteins, within 174 total proteins). Including the additional feature of biofluid label did not significantly resolve this problem (**Figure 2a**), indicating that more expansive biofluid features would be necessary to correct this minor classification discrepancy.

### Feature analysis for importance and correlation with class predictions

During the development of our model, 89 protein features were mined as potentially important to classify these proteins as in vs. out of the nanoparticle corona (**Table S1**). Each feature was examined for the extent of contribution to the overall classification ability of the system using an ANOVA test (**Figure 2b**). This analysis indicates that there is a minimum of approximately ten features to result in sufficient classification ability. We also note that use of approximately 40 features provides a maximum for recall and AUC metrics. If we include up to 89 features, we see a marginal increase in the precision ability of our classifier with a marginal decrease in AUC and recall. As such, experimenters can tune the number of features depending on whether precision or recall is more important: more precise results will be better for experimenters using this tool to correctly identify new nanoparticle-binding proteins, while higher recall results will be better when the opportunity cost of missing a positive corona contributor is more problematic than including a false positive.

Using the feature ranking by ANOVA, the top ten protein features influencing protein adsorption to (GT)_15_-SWCNTs were identified (**Table 1**). Since RFCs do not provide correlational information (i.e., whether a high importance ranking positively or negatively influences protein adsorption), we calculated basic kernel density estimates on distributions of these features and we examined how these distributions changed to hypothesize correlations (**Figure 3**; top ten feature distributions in **Figure S5**). We find that the fraction of solvent-exposed amino acid glycine (normalized to either the total exposed amino acid count or the total amino acid count), the fraction of amino acid glycine, and the fraction of predicted non-secondary structure-associated amino acids correlate positively with the protein being in the corona. Conversely, the fraction of amino acid leucine and the fraction of beta-sheet secondary structure-associated amino acids correlate negatively with being in the corona. Previously, we linearly regressed the log-fold change (ratio of protein amount in the corona vs. in the native biofluid) against physicochemical protein properties to understand protein features that govern corona formation.^52^ In this experimental dataset, high leucine content was similarly determined to be less favorable for protein adsorption to (GT)_15_-SWCNTs. High glycine content was found to be associated with more favorable protein adsorption when included in the regression analysis. However, glycine contribution was not evaluated in the original regression due to correlation with other protein features, as the calculated variance inflation factor was greater than the set threshold value.^52^ As such, glycine content impact could not be deconvoluted from other protein properties. This analysis highlights a benefit of the current RFC over the previously applied linear regression approach, where co-dependent variables must be proactively excluded in the latter case. It should further be noted that secondary structure features were not included in the protein property database for the linear regression analysis due to data sparsity, whereas in the current study we implement BioPython to predict such features from the amino acid sequence without relying on protein structure availability.

**Table 1.** Ordered importance of protein features ranked by ANOVA.

| Ranking | Feature |
| --- | --- |
| 1 | % Amino acid - leucine |
| 2 | % Exposed relative to total exposed amino acids - glycine |
| 3 | % Secondary structure-associated amino acids - non-structure associated |
| 4 | % Exposed relative to total amino acids - glycine |
| 5 | % Amino acid - glycine |
| 6 | % Secondary structure-associated amino acids - sheet |
| 7 | % Exposed relative to total amino acids – tryptophan |
| 8 | % Exposed relative to total amino acids – histidine |
| 9 | % Exposed relative to total exposed amino acids - alanine |
| 10 | % Exposed relative to total exposed amino acids - tryptophan |

**Figure 3.**
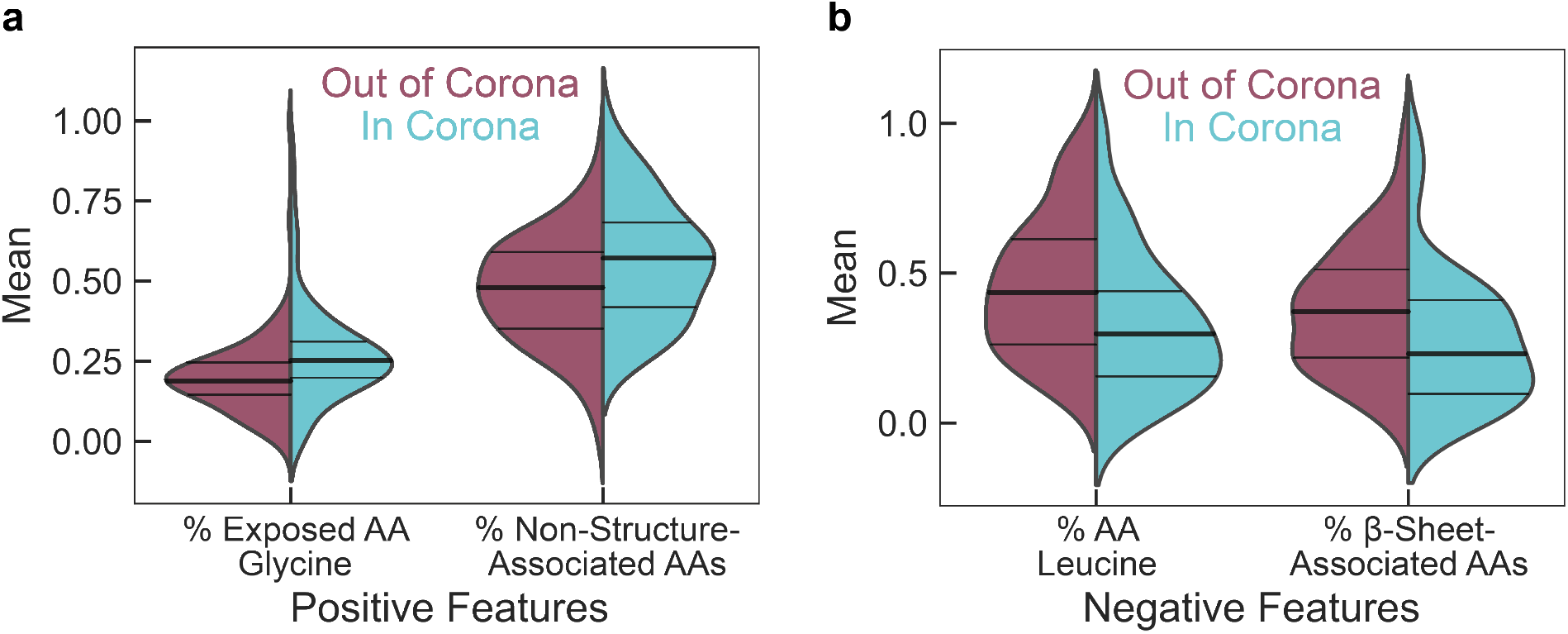
Distribution of the top four normalized feature values for proteins characterized as out of the corona phase (red) vs. in the corona phase (blue) on (GT)_15_-SWCNTs. Protein features that (a) positively influence or (b) negatively influence the probability of a protein being classified as in the corona are denoted by distribution shifts toward 1 or 0, respectively. (a) Positive features include (left) the fraction of solvent-exposed amino acid (AA), glycine, relative to only the solvent-exposed amino acids and (right) the fraction of amino acids not associated with any specific secondary structure motifs. (b) Negative features include (left) the fraction of amino acid, leucine, and (right) the fraction of amino acids associated with a beta-sheet secondary structure.

Our analysis of the top protein features promoting corona binding indicates that more flexible proteins are favorable to bind to (GT)_15_-SWCNTs, as inferred by high glycine content and less strict secondary structural domains. This result is in agreement with previous experimental work demonstrating that peptides and small molecule ligands possessing more conformational flexibility bind more readily to carbon nanotubes.^58,59^ Increased adsorption propensity suggests that more flexible proteins are able to maximize favorable surface contacts with the highly curved SWCNT, in comparison to rigid proteins with energetic penalties associated with adopting new surface-adsorbed conformations. Interestingly, flexibility itself appears in the bottom ten most important protein features for protein corona formation (**Table S2**). This measure of flexibility was calculated by Vihinen *et al*. using normalized, empirically determined B-factors (i.e., Debye–Waller factors) for each residue. B-factors incorporate the dependence on neighboring amino acids with a 9-residue sliding window averaging approach.^60^ With this method, glycine is only the top 8^th^ most flexible residue, posited to be because glycine frequently appears on the protein surface and interior, as well as in tight turns. The restricted mobility of glycine in the interior and turn motifs may reduce the overall flexibility value. As such, our result that high glycine content specifically located on the protein surface is an enriched feature in the corona phase indicates that protein flexibility leads to higher protein corona binding on SWCNTs. In comparison to previous literature, glycine has been found to display a relatively low magnitude, yet still favorable, free energy change upon binding to pristine SWCNTs, as determined by enhanced sampling molecular dynamics.^61^ However, this study was done at the scale of single amino acid analogs. Accordingly, this study disregards the full-protein structural context of each amino acid. Finally, intrinsically disordered proteins have been demonstrated to disperse SWCNTs stably in the aqueous phase even under mild sonication conditions.^62^ Although the non-structure-associated amino acid content that we report is not equivalent to intrinsically disordered domains, our result is in line with these previous experimental findings.

In contrast, our analysis of top protein features that deter corona binding reveals that proteins high in the aliphatic, hydrophobic amino acid leucine and proteins with more planar beta-sheet character are not expected to bind to (GT)_15_-SWCNTs. The finding that hydrophobic leucine does not increase SWCNT binding is not necessarily intuitive, considering that the SWCNT surface is highly hydrophobic. However, this result recapitulates prior literature that nonspecific hydrophobic interactions alone do not drive corona binding;^58,61,63,64^ rather, aromatic hydrophobic amino acids, especially tryptophan, are repeatedly found to be the highest binders to SWCNTs.^58,63–66^ For physical context, (GT)_15_ ssDNA is observed to wrap helically around SWCNTs based on both experiment^67,68^ and modeling,^69,70^ though only covering ~2-25% of the aromatic SWCNT surface.^69,71,72^ Interestingly, the RFC did not highlight aromatic amino acid content (tryptophan, tyrosine, or phenylalanine) as top features for corona binding, although the fraction of exposed tryptophan is the fifth most favorable feature. In studies of isolated amino acids or short peptide sequences, aromatic amino acids seemingly drive adsorption to SWCNTs via π–π interactions with the SWCNT surface. However, in the full protein context, these π–π interactions may not be sufficient to drive initial protein contact with the SWCNT surface, as these hydrophobic amino acids are expected to be predominately buried in the folded protein core. Finally, the finding that high content of amino acids associated with beta-sheet structures leads to low protein adsorption to SWCNTs indicates the difficulty for planar protein secondary structures to adapt to the highly curved nanoparticle surface. This result is in line with previous work demonstrating that the extremely high curvature of carbon nanotubes must be aligned at the amino acid-level of proteins, less the secondary structure level.^58,63^ Overall, the identification of these features is important in helping to predict high biofouling protein types or rationally selecting proteins to bind to nanotubes prior to testing them experimentally.

### Experimental validation of protein binding to SWCNTs

To test the predictive value of our supervised learning model, we applied our classifier to rank a test set of new nanotube-binding proteins and next experimentally tested the expected protein binding order. The classifier was used to predict interaction affinity of over 2,000 total proteins (available for batch download through the UniProt database^53^) with (GT)_15_-SWCNT nanoparticles. Importantly, these proteins represent a broad class of functions and sub-cellular locations, and are distinct from those present in the plasma and CSF training datasets. Protein binding propensity was determined with associated binding probabilities, as summarized in **Table S3**. We then implemented a corona exchange assay to measure real-time, in-solution protein binding dynamics on the nanotube surface, as described previously.^32^ Briefly, the ssDNA originally adsorbed on the SWCNT surface is fluorescently labeled with a Cy5 fluorophore. When near the SWCNT surface, the fluorophore is in a quenched state. Upon addition, proteins will differentially bind to the SWCNT and cause various degrees of ssDNA desorption, as denoted by de-quenching of the Cy5 fluorophore. Thus, fluorescence tracking of the Cy5-ssDNA provides a proxy for protein binding on the SWCNT without requiring fluorescent labeling or other modification of the protein.

The corona exchange assay was used to test a panel of proteins predicted to be in the corona (probability > 0.5) vs. out of the corona (probability < 0.5). Specifically, we tested the protein panel: CD44 antigen and TAR DNA-binding protein 43 (TDP-43) that were predicted to adsorb to (GT)_15_-SWCNTs, vs. transgelin, lysozyme C, ribonuclease pancreatic (RNase A), syntenin-1, L-lactate dehydrogenase A chain (LDH-A), and glutathione S-transferase (GST) that were predicted to not adsorb to (GT)_15_-SWCNTs (classifier results listed in **Table S3**). Protein adsorption based on the end-state fluorescence values matched classifier predicted outcomes of in vs. out of the corona: addition of CD44 antigen and TDP-43 both resulted in sizeable ssDNA desorption from the SWCNT surface, whereas all proteins predicted to be out of the corona produced less ssDNA desorption (**Figure 4a**). However, deviations from exact orderings of predicted outcomes arise within both groups of proteins. For example, the relative ordering of CD44 antigen as the top binding protein followed by TDP-43 is reversed. However, the predicted in-corona probabilities of these two proteins differ by less than 2%. To provide a metric of predicted vs. measured monotonicity, the Spearman’s rank-order correlation coefficient was calculated to be 0.619, in comparison with 0.750 for a previous protein panel comparing DNA desorption end-state vs. proteomic mass spectrometry-derived endstate (**Figure 4c**).^52^ Predicted protein binding probabilities were also compared to rate constants fit to the ssDNA desorption dynamics from the SWCNT surface for each injected protein (kinetic model and fits in SI, **Figure S6**). It is expected that a higher degree of protein binding would correlate with a larger ssDNA desorption rate constant. However, there is poor correlation between the RFC-predicted end-state and experimental dynamics of protein-SWCNT interactions, which may be reconciled with the fact that the RFC was trained on the end-state protein corona rather than the corona composition at earlier time points.

**Figure 4.**
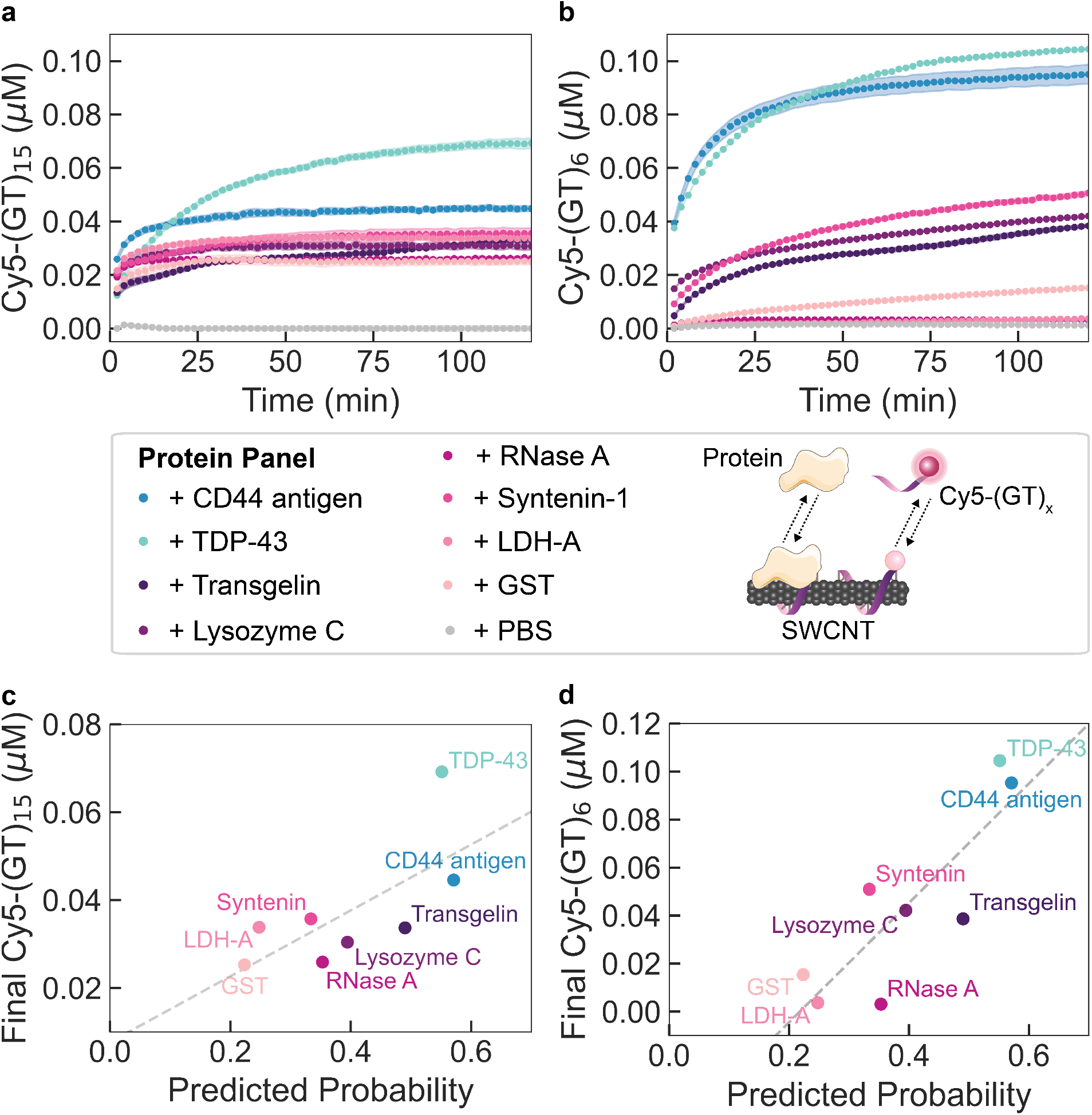
Protein corona dynamics assessed for binding of predicted proteins to (GT)_x_-SWCNTs. (a-b) A corona exchange assay determines binding of a protein panel (each at 80 mg L^-1^ final concentration) to (a) (GT)_15_-SWCNTs or (b) (GT)_6_-SWCNTs (each at 5 mg L^-1^ final concentration). ssDNA desorption from the SWCNT serves as a proxy for protein adsorption. Proteins are predicted by the RFC to be in the corona (probability > 0.5; blue-green colors) or out of the corona (probability < 0.5; purple-pink colors). The protein panel includes: CD44 antigen and TAR DNA-binding protein 43 (TDP-43) (predicted to be in the corona) and transgelin, lysozyme C, ribonuclease pancreatic (RNase A), syntenin-1, L-lactate dehydrogenase A chain (LDH-A), and glutathione S-transferase (GST) (predicted to be out of the corona). Phosphate-buffered saline (PBS) is injected as a control and desorbed ssDNA is normalized to this initial value. Shaded error bars represent standard error between experimental replicates (N = 3). (c-d) End-state desorbed ssDNA is compared to the RFC predicted in-corona probability for (c) (GT)_15_-SWCNTs and (d) (GT)_6_-SWCNTs.

Experimental validation was repeated for the protein panel with Cy5-(GT)_6_-SWCNTs, as this shorter ssDNA oligomer is displaced more readily and thus displays a greater spread in desorption dynamics between protein species (**Figure 4b**).^32^ Moreover, our previous study revealed that the protein corona composition formed on (GT)_6_-SWCNTs was highly similar to that on (GT)_15_-SWCNTs.^52^ The resultant protein panel binding order was largely the same as that of Cy5-(GT)_15_-SWCNTs, with a slightly higher Spearman’s correlation coefficient of 0.667 (**Figure 4d**). These results confirm that the protein binding observed experimentally is mainly driven by the protein interacting directly with the SWCNT nanoparticle surface; the shorter (GT)_6_ ssDNA merely desorbs to a greater extent and thus yields more available SWCNT surface area for protein attachment. This agreement between (GT)_6_- and (GT)_15_-SWCNT datasets further accounts for any mechanistic binding differences in whether protein adsorption causes full or partial ssDNA displacement from the SWCNT surface, where the latter case may occur for longer ssDNA strands.^73^ Comparison of fit rate constants vs. predicted in-corona probabilities reveals a better correlation then that of (GT)_15_, with the exception of RNase A (**Figure S6d**).

Examining the protein identities, it is interesting to note that lysozyme has previously been demonstrated to strongly interact with and disperse pristine carbon nanotubes, in which hydrophobic aromatic amino acids (tryptophan and tyrosine) and cationic amino acids (arginine and lysine) are hypothesized to drive adsorption.^74–78^ Yet, here we find that lysozyme interacts less with pre-dispersed ssDNA-SWCNTs based on the corona exchange results. Therefore, strong lysozyme-SWCNT interaction may hinge upon energetic input employed during the initial SWCNT dispersion process, which likely denatures lysozyme to expose more aromatic residues. This result is important in suggesting that some proteins can only be adsorbed to SWCNT nanoparticles in a partially or fully denatured state, likely compromising their enzymatic activities or protein functions. Another protein of note is CD44, which is overexpressed in cancerous states including upregulation in cancer stem cells.^79^ Toward our goal of facilitating nano-bio construct design, the innate affinity for CD44 to the SWCNT surface could hypothetically be applied to construct a CD44-cell targeted nanotube delivery system.

## Conclusions

In sum, we applied supervised learning methods and developed a classifier to predict protein adsorption on ssDNA-functionalized SWCNTs with 75% accuracy and 68% precision. Ensemble methods performed better in the corona classification task and a random forest classifier scheme was ultimately chosen and optimized. We expand upon prior predictive protein corona work by (i) leveraging quantitative protein corona data,^52^ (ii) redefining corona thresholding, with corresponding prediction probabilities, (iii) establishing a method for classifying proteins based solely on the amino acid sequence of the protein, and (iv) experimentally confirming adsorption in the solution phase with unmodified proteins.^32^ We find that no single nor small group of protein physicochemical features best determine placement in the corona. Rather, over 40 features are useful for protein classification when optimizing all four metrics of accuracy, AUC, precision, and recall. We confirm the need for these protein features by staging them into the classifier feature-by-feature and revalidating our model. Using kernel density estimates, we elucidate protein feature correlation with proteins binding or not binding to SWCNTs. Interestingly, proteins with high solvent-exposed glycine content and more non-structure-associated amino acid content (serving as proxies for protein flexibility) are found to bind in the SWCNT corona, while proteins with high leucine content and beta-sheet-associated amino acid content are not. The classifier then enabled rapid determination of proteins predicted to enter the corona phase from a new protein set, as validated experimentally with a corona exchange assay. Our machine learning algorithm allows us to quicky parse protein properties from a publicly available database to determine protein features of interest for corona formation on SWCNTs.

We intend for this work to support the development of predictive protein corona models that will inform heuristics to rationally select proteins for nanoparticle complexation or to predict biofouling of nanotechnologies. Our model uses amino acid sequence-based prediction of protein corona formation, which could be generalizable across a wide range of bioenvironments. Recent advances in prediction of protein properties from protein sequences alone are promising toward refinement of the protein database we have curated for this classifier, enabling inclusion of biological protein properties that are not reliant on experimental study and manual sequence annotation.^80^ Model accuracy could accordingly be improved by adding structural and geometric protein parameters, such as better-predicted structural motifs, local protein surface curvature, and surface patch hydrophobicity. In the extension of this work, nanoparticle features may be included to enable classification on different nanoparticle types. However, such nanoparticle features should be readily accessible to retain the triviality of classifying new systems. Ultimately, *in silico* protein corona prediction will support the design of nanotechnologies that can be more seamlessly implemented in biological systems with reduced need for experimental mass spectrometry-based proteomic characterization and analysis. The ability to predict adsorption of specific proteins will enable connection to downstream cellular responses, toxicity outcomes, and overall nanotechnology functionality. The developed classifier provides a preliminary tool for both predicting key proteins expected to take part in biofouling and rapid prescreening of protein candidates in rationally designed nanobiotechnologies.

## Methods

### Database development

Protein information was downloaded from UniProt,^53^ including amino acid sequences (FASTA format) and sequence annotations. Amino acid sequences were used to generate a series of physicochemical protein properties using BioPython’s Protein Analysis module (**Table S1**).^54^ Amino acid sequences were additionally analyzed by NetSurfP 2.0^55^ to determine solvent accessibility, including relative solvent accessibility (RSA), absolute solvent accessibility (ASA), and fractions of each amino acid exposed surface area relative to either all amino acids or only other exposed amino acid surface area. To collate this data, we programmatically created submissions from UniProt protein sequence entries to NetSurfP 2.0, aligning with our goal of creating an easily expandable database. The resulting data was processed and merged with the BioPython analysis. The complete database was run normalized with a Min Max Scalar from Scikit-Learn^81^ before being subset and fit to the classification model. Code for this and all subsequent sections can be found in the GitHub link provided.

### Criteria for in-corona placement

Using the method described previously for protein corona studies by LC-MS/MS,^52^ quantitative data was obtained for proteins adsorbing to (GT)_15_-SWCNTs in two different human biofluids: blood plasma and cerebrospinal fluid. First, proteins with abundances (*A_corona_*) greater than the control of protein abundances in biofluids alone (*A_biofluid_*) were assigned as in the corona (i.e., enriched in the corona relative to the biofluid). Second, an exponential decay, *n* = *n*_0_ exp(–*kA*), was fit to the distribution of abundances for the remaining proteins, where *n*_0_ and *k* are fitting parameters. An abundance threshold (*A_threshold_*) was selected at a value where the exponential decay fell to a value of *n*_0_exp (–*p*), or *A_threshold_* = *p/k*, where *p* is an optimization parameter. Proteins with an abundance greater than *A_threshold_* were assigned as being in the corona. We varied *p* between 0 and 3.5 and chose the value 2.25, which optimized the performance of the classifier following training (**Figure 1a**) and was used for the remainder of the analysis. Corona thresholding was originally completed with Otsu’s method, a technique generally implemented for image thresholding.^82^ However, employing Otsu’s method resulted in only 3-5 proteins placed in the corona for each biofluid. Although the classifier was highly accurate at identifying these proteins, the number of proteins selected was not fully representative of the corona and we accordingly implemented our modified thresholding method described above.

### Classifier selection

The use of a random forest classifier (RFC), logistic regression, bagging classifier, gradient boosting classifier, AdaBoost classifier, and XGBoost classifier were evaluated. The RFC, logistic regression, bagging classifier, gradient boosting classifier, and AdaBoost classifier were imported from Scikit-Learn.^81^ The XGBoost classifier was imported from XGBoost^83^ for use with Scikit-Learn. AdaBoost and bagging classifiers were tested with an underlying support vector machine, decision tree, and logistic regression. The gradient boosting classifier was tested with an underlying decision tree. XGBoost was tested with an underlying decision tree as well as 100 parallel trees.

The RFC performed best and was accordingly chosen for the remainder of the work. The classifier was next validated using a stratified shuffle split (100 repeats) validation to ensure consistent levels of the minority class. The minority class here is the in-corona class which has less proteins than the out-of-corona class. The shuffle split retained 10% of the dataset for corona validation. The training split was augmented with entries developed from SMOTE, as detailed in the main text. Results were collected for each fold. For cross-biofluid tests, the percentage of proteins in the test set was varied to keep the same number of proteins in the test set equal to 10% of the total number of proteins used for mixed biofluid cases. The adjusted value was set by scaling 10% by a factor of the total number of proteins divided by the number of proteins in the test biofluid (Plasma: 1.55; CSF: 2.81).

### Hyperparameter tuning

Using Scikit-Learn’s GridSearchCV,^81^ a wide range of hyperparameters, such as number or depth of trees, were tested with the classifier. With each set of hyperparameters the model was validated using the method dictated in the **Classifier Selection** section and scored. The classifier was chosen with the hyperparameters optimized for precision using GridSeachCV.

### Dimensionality reduction

To understand the effects of each feature (i.e. variable describing the protein of interest) on the total system, features were ranked using Scikit-Learn’s SelectKBest function.^81^ Using the ranking established from SelectKBest, the database features were unmasked one-by-one running the classifier as described in the **Classifier Selection** section until all features had been added in. Metric results were saved, and statistics were calculated.

### New prediction targets

The classifier was tested against a list of 996 cytoplasmic proteins and 999 nuclear proteins (available for batch download through the UniProt database^53^), together with 45 readily accessible proteins or proteins of interest for SWCNT-based sensing and delivery applications. Amino acid sequences for these proteins were downloaded from UniProt and processed through the database development workflow described above. This new complete protein database was then processed through the classifier k+1 times. The first k times were completed through the described k-fold validation using the combined datasets for (GT)_15_-SWCNTs in plasma and cerebrospinal fluid as the training and verification data. Predictions were recorded at the end of each fold. The last time new proteins were run, all available data was used to train the classifier; this last classifier then provided predictions on the new proteins, as listed in **Table S3**. Training the classifier exclusively on the (GT)_6_-SWCNT in plasma LC-MS/MS dataset revealed qualitatively similar results.

### Synthesis of ssDNA-SWCNTs

Suspensions of single-walled carbon nanotube (SWCNTs) with fluorophore-labeled single-stranded DNA (Cy5-(GT)_15_ or Cy5-(GT)_6_) were prepared with 0.2 mg of mixed-chirality SWCNTs (small diameter HiPco^™^ SWCNTs, NanoIntegris) and 20 μM of ssDNA (3’ Cy5-labeled custom ssDNA oligos with HPLC purification, Integrated DNA Technologies, Inc.; excitation 648 nm, emission 668 nm) added in 1 mL total volume of 0.1X phosphate-buffered saline (PBS; note 1X is 137 mM NaCl, 2.7 mM KCl, 10 mM Na_2_HPO_4_, 1.8 mM KH_2_PO_4_).^32^ This mixture was probe-tip sonicated for 10 min in an ice bath (3 mm probe tip at 50% amplitude, 5-6 W, Cole-Parmer Ultrasonic Processor). Cy5-ssDNA-SWCNT suspensions were centrifuged to pellet insoluble SWCNT bundles and contaminants (16.1 krcf, 30 min). The supernatant containing product was collected and Cy5-ssDNA-SWCNT concentration was calculated with measured sample absorbance at 910 nm (NanoDrop One, Thermo Scientific) and the empirical extinction coefficient, ε_910nm_=0.02554 L mg^-1^ cm^-1^.^84^ Cy5-ssDNA-SWCNTs were stored at 4°C until use, at which point the solution was diluted to a working concentration of 10 mg L^-1^ in 1X PBS ≥ 2 h prior to use.

### Preparation of proteins

Proteins were sourced as listed in **Table 2**. Lyophilized proteins were reconstituted to the listed concentration in PBS, tilting intermittently to dissolve for 15 min, and filtering with 0.2 μm syringe filters (cellulose acetate membrane, VWR International). All proteins were purified with desalting columns (Zeba Spin Desalting Columns, 0.5 mL with 7 kDa MWCO, Thermo Fisher Scientific) by washing with PBS three times (centrifuging 1500 rcf, 1 min), centrifuging with sample (1500 rcf, 2 min), and retaining sample in flow-through solution. Resulting protein concentration was measured with the Qubit Protein Assay (Thermo Fisher Scientific).

**Table 2.** Purchased protein specifications.

| Protein | Manufacturer | Catalog # | Lot # | Source | Notes |
| --- | --- | --- | --- | --- | --- |
| CD44 antigen | Acro Biosystems | CD4-H5226 | 616-784F1-G8 | Human, expressed in HEK293 | 6X His tag; >95% purity |
| TAR DNA-binding protein 43 (TDP-43) | R&D Systems | AP-190 | 22675420A | Recombinant human, expressed in E. coli | >85% purity |
| Transgelin (TAGLN) | MyBioSource | MBS144070 | 1011PTAGLN30 | Recombinant human, expressed in E. coli | 20X His tag; >85% purity |
| Lysozyme C | Sigma | L2879 | SLCF2361 | From chicken egg white | ≥80% purity |
| Ribonuclease pancreatic (RNase A) | New England BioLabs | T3018L |  | Purified from cow pancreas |  |
| Syntenin-1 | Novus Biologicals | NBP1-50893 | 1082301 | Recombinant human, expressed in E. coli | 6X His tag; >90% purity |
| L-lactate dehydrogenase A chain (LDH-A) | Sigma-Aldrich | 10127230001 | 42032824 | From rabbit muscle |  |
| Glutathione S-transferase (GST) | Abcam | ab86775 | GR3377596-1 | Recombinant mouse, expressed in E. coli | >95% purity |

### Corona exchange assay

Corona dynamics were measured as described previously.^32^ Briefly, equal volumes (25 μL) of ssDNA-Cy5-SWCNT and FAM-protein at 2X working concentration were added via multichannel pipette into a 96-well PCR plate (Bio-Rad) and mixed by pipetting. The PCR plate was sealed with an optically transparent adhesive seal (Bio-Rad) and briefly spun down on a benchtop centrifuge. Fluorescence was measured as a function of time using a Bio-Rad CFX96 Real Time qPCR System, scanning all manufacturer set color channels (FAM, HEX, Texas Red, Cy5, Quasar 705) every 30 s at 22.5 °C, with lid heating off. Fluorescence time series were analyzed without default background correction. Of note, fluorophore dequenching indicates that the 3’ end of the Cy5-tagged ssDNA was displaced from the SWCNT surface and may not indicate complete ssDNA strand desorption.

## Acknowledgements

We acknowledge support of the IGI LGR ERA, GlaxoSmithKline, and Citris/Banatao Seed Funding. We acknowledge support of a Burroughs Wellcome Fund Career Award at the Scientific Interface (CASI) (to M.P.L.), a Dreyfus foundation award (to M.P.L.), a Stanley Fahn PDF Junior Faculty Grant with Award # PF-JFA-1760 (to M.P.L.), a Beckman Foundation Young Investigator Award (to M.P.L.), an NIH MIRA award (to M.P.L.), an NSF CAREER award (to M.P.L), an NSF CBET award (to M.P.L.), an NSF CGEM award (to M.P.L.), a FFAR Young Investigator award (to M.P.L.), a CZI investigator award (to M.P.L), a Sloan Foundation Award (to M.P.L.), a USDA BBT EAGER award (to M.P.L), a USDA NIFA Award (to M.P.L), a Moore Foundation Award (to M.P.L.), a Cisco Research Center grant (to M.P.L), and a DARPA Young Investigator Award (to M.P.L.). M.P.L. is a Chan Zuckerberg Biohub investigator, a Hellen Wills Neuroscience Institute Investigator, and an IGI Investigator. N.O., R.L.P., and J.W. acknowledge the support of NSF Graduate Research Fellowships (NSF DGE 1752814). J.T.D.B.-O. acknowledges the support of an Early Investigator Research Award from the Congressionally Directed Medical Research Program through the U.S. Department of Defense. We would like to acknowledge the use of medical clipart from Servier Medical Art by Servier (http://smart.servier.com), licensed under a Creative Commons Attribution 3.0 Unported License.

