## Supplementary Information for "Supervised Learning Model Predicts Protein Adsorption to Carbon Nanotubes"

**Table S1**. Protein property list.

| Protein Property | Calculated By | Implementation in Code |
|---|---|---|
| % Amino Acid - Alanine (A) | BioPython | frac_aa_A |
| % Amino Acid - Cysteine (C) | BioPython | frac_aa_C |
| % Amino Acid - Aspartic Acid (D) | BioPython | frac_aa_D |
| % Amino Acid - Glutamic Acid (E) | BioPython | frac_aa_E |
| % Amino Acid - Phenylalanine (F) | BioPython | frac_aa_F |
| % Amino Acid - Glycine (G) | BioPython | frac_aa_G |
| % Amino Acid - Histidine (H) | BioPython | frac_aa_H |
| % Amino Acid - Isoleucine (I) | BioPython | frac_aa_I |
| % Amino Acid - Lysine (K) | BioPython | frac_aa_K |
| % Amino Acid - Leucine (L) | BioPython | frac_aa_L |
| % Amino Acid - Methionine (M) | BioPython | frac_aa_M |
| % Amino Acid - Asparagine (N) | BioPython | frac_aa_N |
| % Amino Acid - Proline (P) | BioPython | frac_aa_P |
| % Amino Acid - Glutamine (Q) | BioPython | frac_aa_Q |
| % Amino Acid - Arginine (R) | BioPython | frac_aa_R |
| % Amino Acid - Serine (S) | BioPython | frac_aa_S |
| % Amino Acid - Threonine (T) | BioPython | frac_aa_T |
| % Amino Acid - Valine (V) | BioPython | frac_aa_V |
| % Amino Acid - Tryptophan (W) | BioPython | frac_aa_W |
| % Amino Acid - Tyrosine (Y) | BioPython | frac_aa_Y |
| Molecular Weight | BioPython | molecular_weight |
| Aromaticity | BioPython | aromaticity |
| Instability Index | BioPython | instability_index |
| Flexibility - Mean | BioPython | flexibility_mean |
| Flexibility - Standard Deviation | BioPython | flexibility_std |
| Flexibility - Variance | BioPython | flexibility_var |
| Flexibility - Max | BioPython | flexibility_max |
| Flexibility - Min | BioPython | flexibility_min |
| Flexibility - Median | BioPython | flexibility_median |
| Isoelectric Point | BioPython | isoelectric_point |
| % Secondary Structure-Associated Amino Acids – Helix (V, I, Y, F, W, L) | BioPython | secondary_structure_fraction_helix |
| % Secondary Structure-Associated Amino Acids – Turn (N, P, G, S) | BioPython | secondary_structure_fraction_turn |
| % Secondary Structure-Associated Amino Acids – Sheet (E, M, A, L) | BioPython | secondary_structure_fraction_sheet |
| % Secondary Structure-Associated Amino Acids - Non-Structure Associated | BioPython | secondary_structure_fraction_disordered |
| Length | BioPython | length |
| Mass | BioPython | mass |
| % Amino Acids Exposed | NetSurfP | fraction_exposed |
| % Amino Acids Buried | NetSurfP | fraction_buried |
| % Exposed Nonpolar Amino Acids / Total Amino Acids | NetSurfP | fraction_exposed_nonpolar_total |
| % Exposed Nonpolar Amino Acids / Exposed | NetSurfP | fraction_exposed_nonpolar_exposed |

| | | |
|---|---|---|
| Relative Surface Area (RSA) - Mean | NetSurfP | rsa_mean |
| Relative Surface Area - Median | NetSurfP | rsa_median |
| Relative Surface Area - Standard Deviation | NetSurfP | rsa_std |
| Absolute Surface Area (ASA) - Sum | NetSurfP | asa_sum |
| % Exposed Amino Acid A / Total Amino Acids | NetSurfP | fraction_total_exposed_A |
| % Exposed Amino Acid C / Total Amino Acids | NetSurfP | fraction_total_exposed_C |
| % Exposed Amino Acid D / Total Amino Acids | NetSurfP | fraction_total_exposed_D |
| % Exposed Amino Acid E / Total Amino Acids | NetSurfP | fraction_total_exposed_E |
| % Exposed Amino Acid F / Total Amino Acids | NetSurfP | fraction_total_exposed_F |
| % Exposed Amino Acid G / Total Amino Acids | NetSurfP | fraction_total_exposed_G |
| % Exposed Amino Acid H / Total Amino Acids | NetSurfP | fraction_total_exposed_H |
| % Exposed Amino Acid I / Total Amino Acids | NetSurfP | fraction_total_exposed_I |
| % Exposed Amino Acid K / Total Amino Acids | NetSurfP | fraction_total_exposed_K |
| % Exposed Amino Acid L / Total Amino Acids | NetSurfP | fraction_total_exposed_L |
| % Exposed Amino Acid M / Total Amino Acids | NetSurfP | fraction_total_exposed_M |
| % Exposed Amino Acid N / Total Amino Acids | NetSurfP | fraction_total_exposed_N |
| % Exposed Amino Acid P / Total Amino Acids | NetSurfP | fraction_total_exposed_P |
| % Exposed Amino Acid Q / Total Amino Acids | NetSurfP | fraction_total_exposed_Q |
| % Exposed Amino Acid R / Total Amino Acids | NetSurfP | fraction_total_exposed_R |
| % Exposed Amino Acid S / Total Amino Acids | NetSurfP | fraction_total_exposed_S |
| % Exposed Amino Acid T / Total Amino Acids | NetSurfP | fraction_total_exposed_T |
| % Exposed Amino Acid V / Total Amino Acids | NetSurfP | fraction_total_exposed_V |
| % Exposed Amino Acid W / Total Amino Acids | NetSurfP | fraction_total_exposed_W |
| % Exposed Amino Acid Y / Total Amino Acids | NetSurfP | fraction_total_exposed_Y |
| % Exposed Amino Acid A / Total Exposed | NetSurfP | fraction_exposed_exposed_A |
| % Exposed Amino Acid C / Total Exposed | NetSurfP | fraction_exposed_exposed_C |
| % Exposed Amino Acid D / Total Exposed | NetSurfP | fraction_exposed_exposed_D |
| % Exposed Amino Acid E / Total Exposed | NetSurfP | fraction_exposed_exposed_E |
| % Exposed Amino Acid F / Total Exposed | NetSurfP | fraction_exposed_exposed_F |
| % Exposed Amino Acid G / Total Exposed | NetSurfP | fraction_exposed_exposed_G |
| % Exposed Amino Acid H / Total Exposed | NetSurfP | fraction_exposed_exposed_H |
| % Exposed Amino Acid I / Total Exposed | NetSurfP | fraction_exposed_exposed_I |
| % Exposed Amino Acid K / Total Exposed | NetSurfP | fraction_exposed_exposed_K |
| % Exposed Amino Acid L / Total Exposed | NetSurfP | fraction_exposed_exposed_L |
| % Exposed Amino Acid M / Total Exposed | NetSurfP | fraction_exposed_exposed_M |
| % Exposed Amino Acid N / Total Exposed | NetSurfP | fraction_exposed_exposed_N |
| % Exposed Amino Acid P / Total Exposed | NetSurfP | fraction_exposed_exposed_P |
| % Exposed Amino Acid Q / Total Exposed | NetSurfP | fraction_exposed_exposed_Q |
| % Exposed Amino Acid R / Total Exposed | NetSurfP | fraction_exposed_exposed_R |
| % Exposed Amino Acid S / Total Exposed | NetSurfP | fraction_exposed_exposed_S |
| % Exposed Amino Acid T / Total Exposed | NetSurfP | fraction_exposed_exposed_T |
| % Exposed Amino Acid V / Total Exposed | NetSurfP | fraction_exposed_exposed_V |
| % Exposed Amino Acid W / Total Exposed | NetSurfP | fraction_exposed_exposed_W |
| % Exposed Amino Acid Y / Total Exposed | NetSurfP | fraction_exposed_exposed_Y |

| | | |
|---|---|---|
| Sum of Absolute Surface Area / Total Mass | NetSurfP | asa_sum_normalized |
| % Secondary Structure - Helix | NetSurfP | nsp_secondary_structure_helix |
| % Secondary Structure - Sheet | NetSurfP | nsp_secondary_structure_sheet |
| % Secondary Structure - Coil | NetSurfP | nsp_secondary_structure_coiled |
| % Secondary Structure - Disordered | NetSurfP | nsp_disordered |

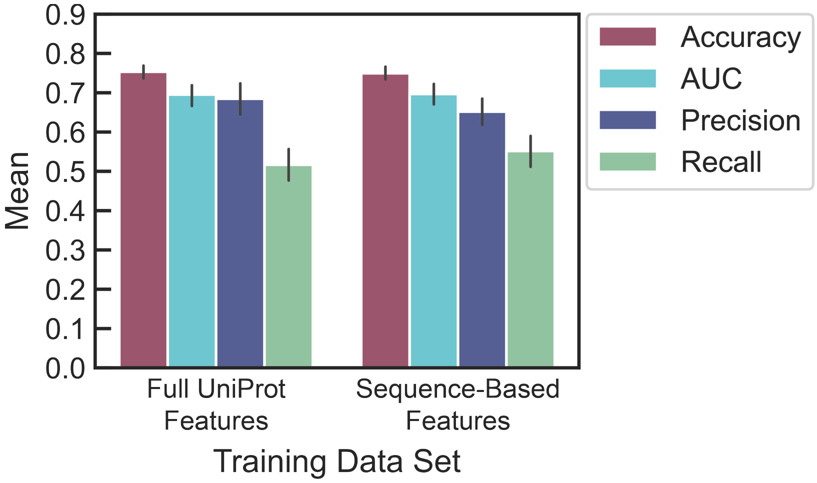


**Figure S1**. Comparison of classifier performance with all biological and physicochemical features available through UniProt (left) and with only amino acid-sequence derived features as listed in **Table S1** (right).

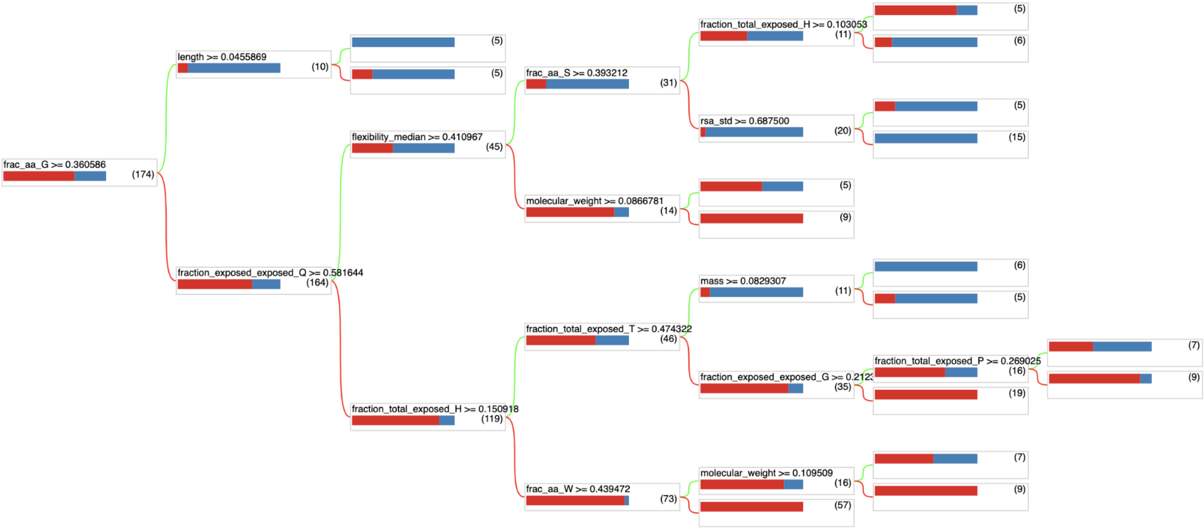


**Figure S2**. Example random forest tree (1 of 300 trees). Each tree is traversed left to right. If the node criteria evaluates to true, the green path is taken. If the node criteria evaluates to false, the red path is taken. This process is repeated until the protein reaches a terminal node then a vote is recorded. Total votes among all trees are tallied and a decision is made by majority rules. Bars at each node indicate the percentage of data at that node that is in each phase (blue - in corona, red - out of corona). Parentheses indicate the number of data points at each node. Created using TensorFlow-Decision Forests.[85]

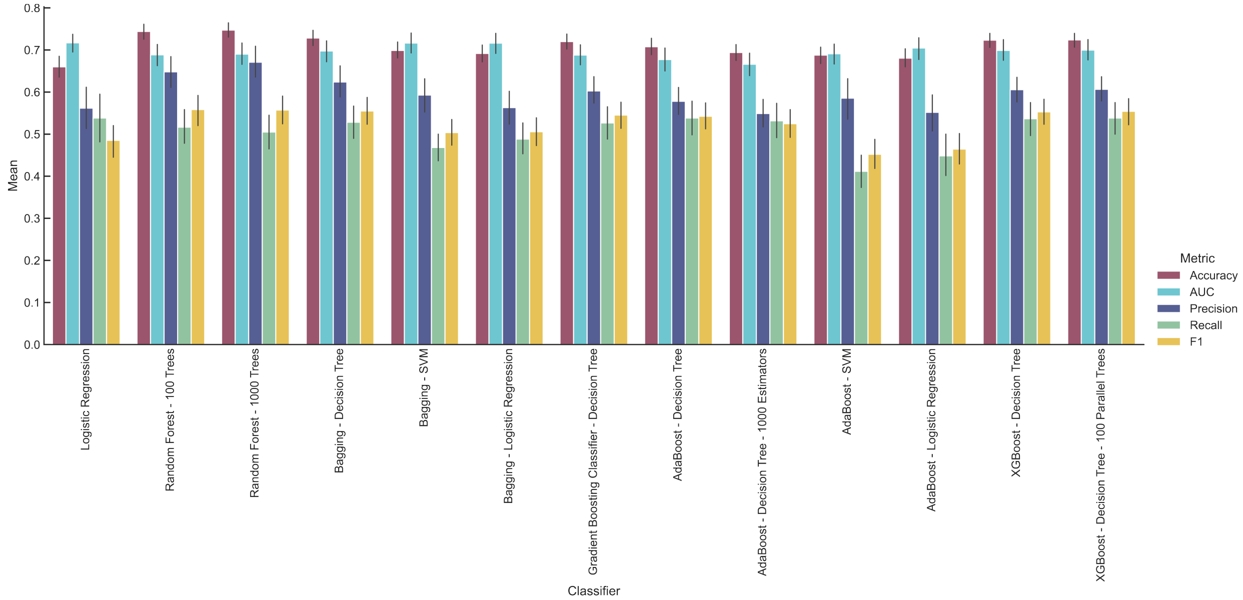


**Figure S3**. Comparison of output metrics for a panel of classifiers. Classifiers tested: logistic regression, random forest (100 or 1000 trees), bagging (decision tree, SVM, logistic regression), gradient boosting classifier (decision tree), AdaBoost (decision tree and with 1000 estimators, SVM, logistic regression), and XGBoost (decision tree and with 100 parallel trees). Corresponding classifier metrics are reported for each: accuracy, area under the receiver operating curve (AUC), precision, and recall.

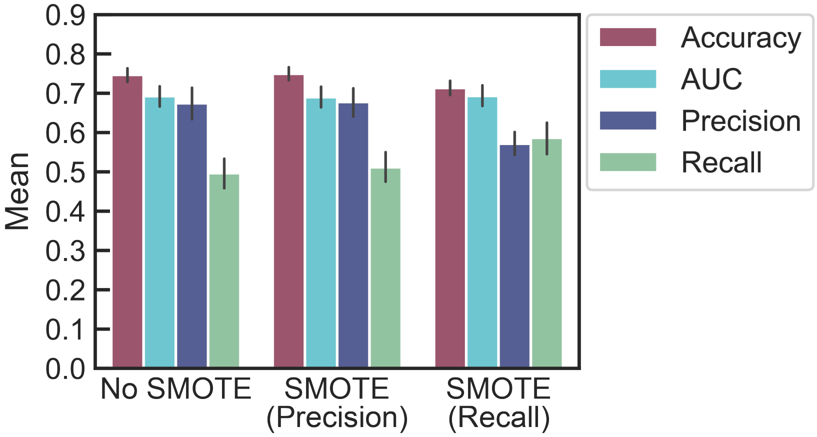


**Figure S4**. Comparison of classifier performance prior to (left) and upon introducing a synthetic minority over-sampling technique (SMOTE), optimized for either precision (middle) with a SMOTE ratio of 0.5 or recall (right) with a SMOTE ratio of 1.

**a**

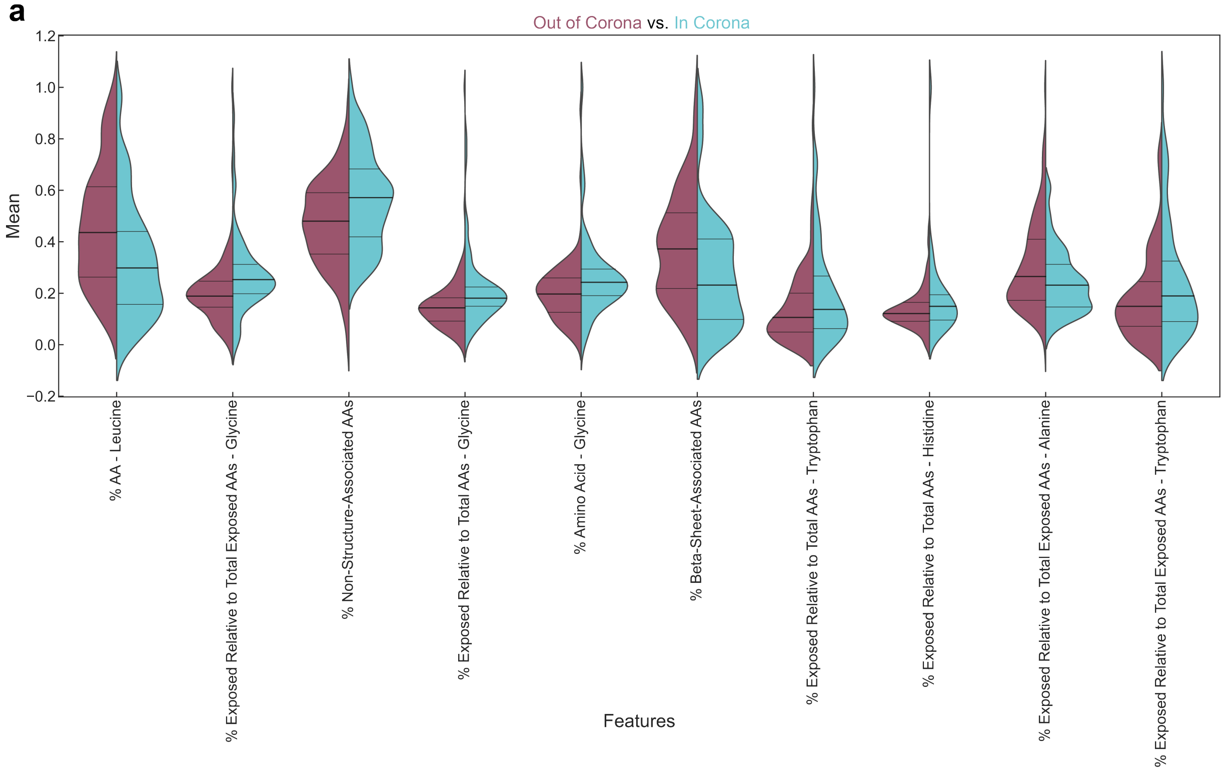


**b**

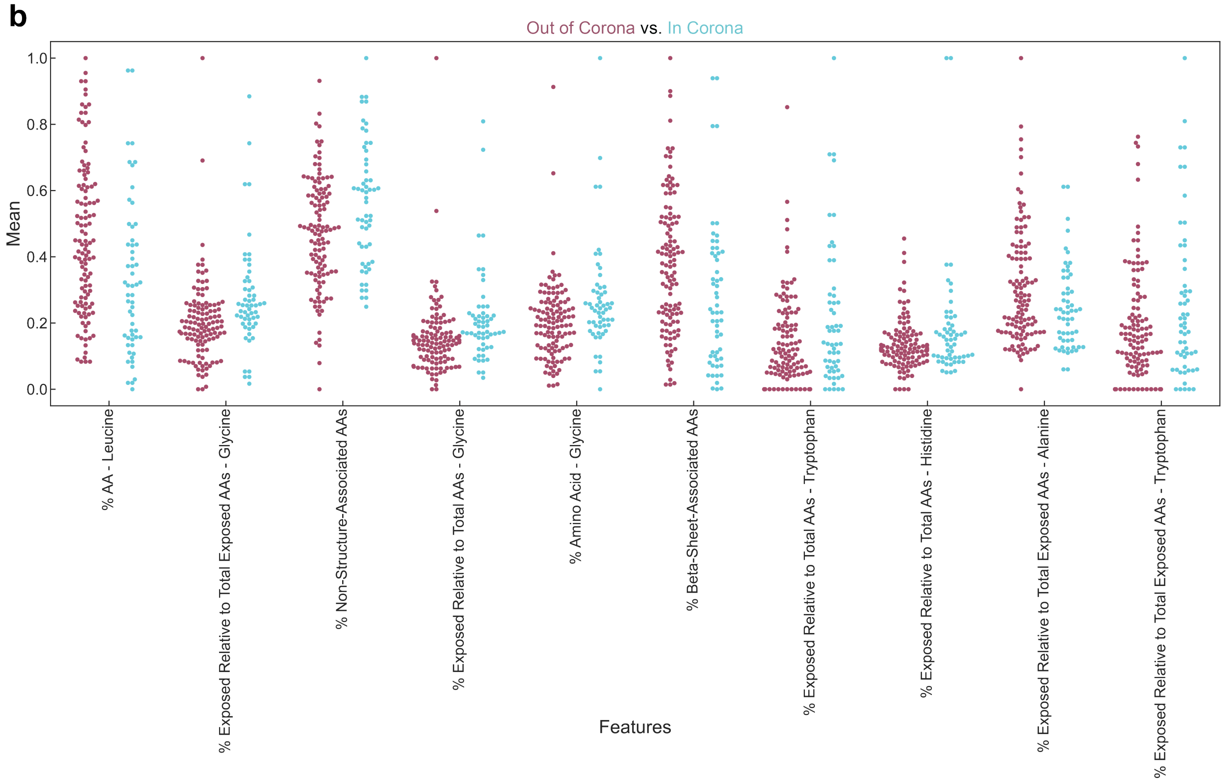


**Figure S5**. Distribution of the top ten normalized feature values for proteins characterized as out of the corona phase (red) vs. in the corona phase (blue) on $(GT)_{15}$-SWCNTs. Protein features that positively influence or negatively influence the probability of a protein being classified as in the corona are denoted by distribution shifts toward 1 or 0, respectively. Protein features are represented with (a) violin plots and (b) scatter distribution plots.

**Table S2**. Ordered importance of protein features by weight.

| Ranking | Top Features | Weight |
|---|---|---|
| 1 | % Exposed relative to total exposed amino acids - glycine | 0.03288 |
| 2 | % Exposed relative to total amino acids - glycine | 0.03189 |
| 3 | % Secondary structure-associated amino acids - sheet | 0.02714 |
| 4 | % Amino acid - leucine | 0.02481 |
| 5 | Instability index | 0.01978 |
| 6 | % Amino acid - glycine | 0.01915 |
| 7 | % Secondary structure-associated amino acids - non-structure associated | 0.01890 |
| 8 | % Amino acid - alanine | 0.01835 |
| 9 | % Amino acid - tryptophan | 0.01664 |
| 10 | % Exposed relative to total amino acids - histidine | 0.01407 |
| **Ranking** | **Bottom Features** | **Weight** |
| 1 | % Exposed relative to total exposed amino acids - valine | 0.00713 |
| 2 | % Amino acid – aspartic acid | 0.00713 |
| 3 | Flexibility - mean | 0.00709 |
| 4 | % Exposed relative to total amino acids - aspartic acid | 0.00687 |
| 5 | % Exposed relative to total amino acids - methionine | 0.00666 |
| 6 | % Exposed relative to total exposed amino acids - glutamic acid | 0.00648 |
| 7 | Molecular weight | 0.00635 |
| 8 | % Secondary structure - sheet | 0.00634 |
| 9 | % Exposed relative to total exposed amino acids - cysteine | 0.00573 |
| 10 | Sequence length | 0.00545 |

**Table S3**. Classifier predictions of high- vs. low-binding proteins on $(GT)_{15}$-SWCNTs.

| Ranking | Protein | Accession Number | Probability |
|---|---|---|---|
| In 1 | CD44 antigen | P16070 | 57.10% |
| In 2 | TAR DNA-binding protein 43 (TDP-43) | Q13148 | 55.14% |
| Out 1 | Transgelin | Q01995 | 49.02% |
| Out 2 | Lysozyme C | P00698 | 39.46% |
| Out 3 | Ribonuclease pancreatic (RNase A) | P07998 | 35.31% |
| Out 4 | Syntenin-1 | O00560 | 33.38% |
| Out 5 | L-lactate dehydrogenase A chain (LDH-A) | P00338 | 24.77% |
| Out 6 | Glutathione S-transferase (GST) | Q8MU52 | 22.35% |

A simple kinetic model was fit to the ssDNA desorption data from the corona exchange assay:

$$D + * \underset{k_2}{\overset{k_1}{\rightleftarrows}} D *$$

$$[*]_{\mathrm{tot}} = [*] + [D *]$$

**Equation 1**

where *D* is ssDNA, * is a SWCNT surface site, and *D** is ssDNA bound to a SWCNT surface site. Square brackets represent concentrations. Rate constants $k_1$, $k_2$, and total concentration of SWCNT surface sites $[*]_{\mathrm{tot}}$ were fit using an ordinary least-squares regression (**Figure S6**).

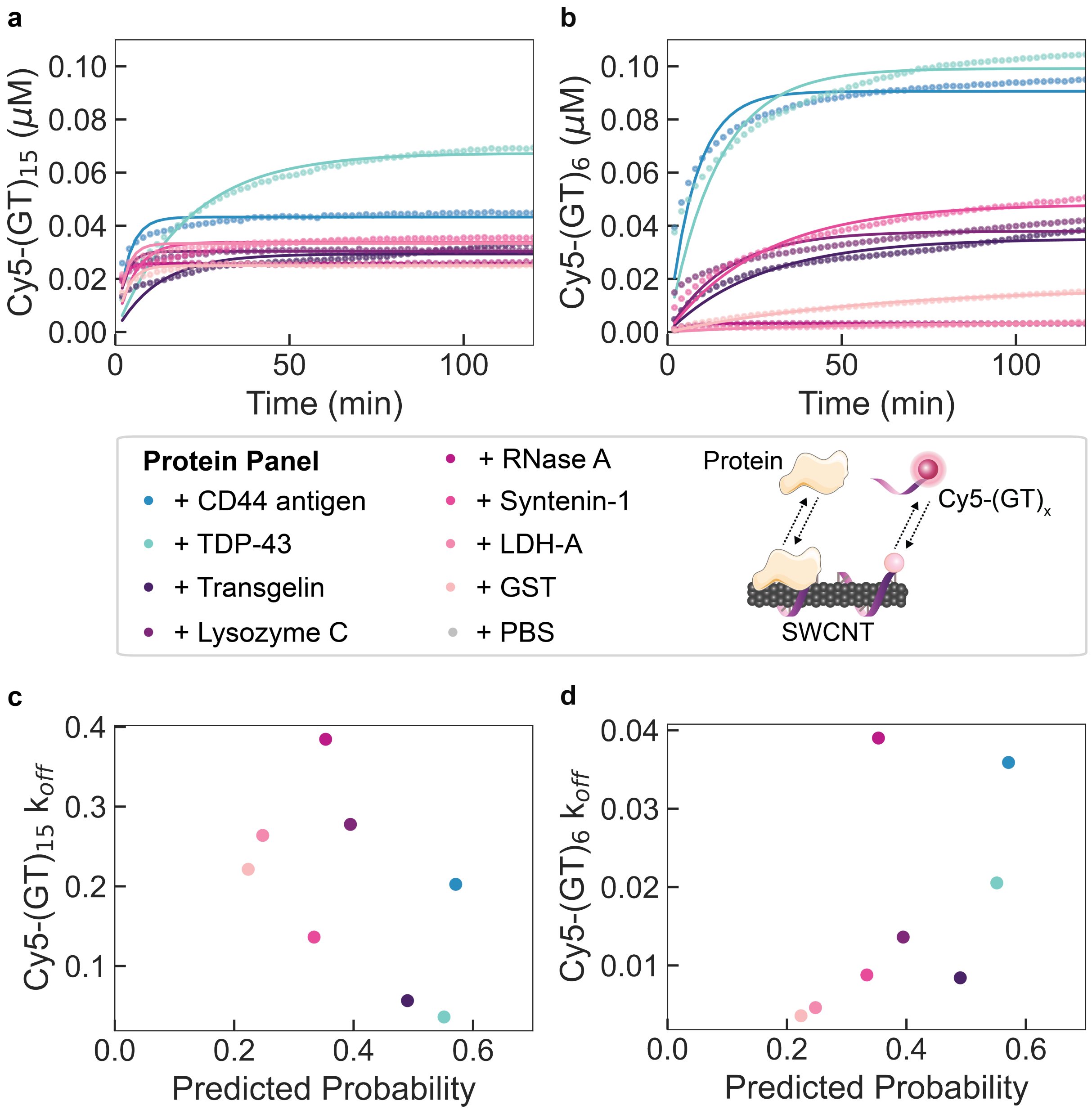


**Figure S6**. Protein corona dynamics assessed for binding of predicted proteins to $(GT)_x$-SWCNTs. (a-b) A corona exchange assay determines binding of a protein panel (each at 80 mg $L^{-1}$ final concentration) to (a) $(GT)_{15}$-SWCNTs or (b) $(GT)_6$-SWCNTs (each at 5 mg $L^{-1}$ final concentration). ssDNA desorption from the SWCNT serves as a proxy for protein adsorption. Proteins are predicted by the RFC to be in the corona (probability > 0.5; blue-green colors) or out of the corona (probability < 0.5; purple-pink colors). The protein panel includes: CD44 antigen and TAR DNA-binding protein 43 (TDP-43) (predicted to be in the corona) and transgelin, lysozyme C, ribonuclease pancreatic (RNase A), syntenin-1, L-lactate dehydrogenase A chain (LDH-A), and glutathione S-transferase (GST) (predicted to be out of the corona). Phosphate-buffered saline (PBS) is injected as a control and desorbed ssDNA is normalized to this initial value. Shaded error bars represent standard error between experimental replicates (N = 3). (c-d) Rate constants of ssDNA desorption are compared to the RFC predicted in-corona probability for (c) $(GT)_{15}$-SWCNTs and (d) $(GT)_6$-SWCNTs.
